# B cells imprinted by ancestral SARS-CoV-2 develop pan-sarbecovirus neutralization in immune recalls

**DOI:** 10.1101/2024.10.13.618110

**Authors:** Xixian Chen, Ling Li, Ruiping Du, Zuowei Wang, Yunjian Li, Yi Sun, Rongrong Qin, Hualong Feng, Lin Hu, Xuanyi Chen, Maosheng Lu, Xueyan Huang, Liwei Jiang, Teng Zuo

**Affiliations:** Laboratory of Immunoengineering, Institute of Health and Medical Technology, Hefei Institutes of Physical Science, Chinese Academy of Sciences, Hefei 230031, China; University of Science and Technology of China, Hefei 230026, China

## Abstract

A key question on ancestral SARS-CoV-2 immune imprinting is to what extent imprinted B cells can develop neutralizing breadth and potency in immune recalls. Here, we longitudinally tracked B cells recognizing wild-type spike in two individuals, who were sequentially infected by Omicron variants after receiving mRNA vaccines. Functional and genetic analysis of 632 monoclonal antibodies (mAbs) from those B cells reveals that mAbs cloned after second infection have dramatically enhanced neutralizing breadth and potency, which is attributed to recall and maturation of pre-existing memory B cells. Among the 11 mAbs that potently neutralize SARS-CoV-2 variants from wild-type to KP.3, 5 mAbs are classified into public clonotypes encoded by IGHV3-53 or IGHV3-66, whereas the rest belong to a rarely reported clonotype encoded by IGHV3-74. Notably, IGHV3-74 mAbs can also broadly neutralize other sarbecoviruses by targeting a novel epitope on receptor-binding domain of spike. These results support that ancestral SARS-CoV-2 immune imprinting can be harnessed in developing pan-SARS-CoV-2 and even pan-sarbecovirus vaccines.

**Summary:** Chen et al. demonstrate that B cells imprinted by ancestral SARS-CoV-2 have tremendous potential to develop neutralizing breadth and potency in repeated immune recalls driven by Omicron variants, implicating that ancestral SARS-CoV-2 immune imprinting can be harnessed in developing pan-SARS-CoV-2 and even pan-sarbecovirus vaccines.

## Introduction

Adaptive immune system adopts multiple mechanisms to ensure specific and diverse antibody responses against various pathogens. During B cell development, V(D)J recombination generates a vast diversity of B cell antigen receptors (BCRs) for naïve B cells (Tonegawa, 1983). When infection or vaccination occurs, a small portion of naïve B cell repertoire get activated as their BCRs recognize corresponding antigens by chance (Batista and Harwood, 2009). Activated B cells either differentiate into short-lived plasmablasts (PBs) to produce the initial wave of antibodies, or enter germinal centers (GCs) to undergo repeated rounds of somatic hypermutation and affinity-dependent selection, a process known as antibody affinity maturation (Victora and Nussenzweig, 2022). The evolved GC B cells eventually differentiate into long-lived plasma cells (PCs) to constantly secrete antibodies for months to years, or memory B cells to wait for future encounter with the same or similar antigens (Victora and Nussenzweig, 2022). Compared with primary antigen exposure, secondary antigen exposure typically leads to much faster and stronger antibody responses as recalled memory B cells rapidly differentiate into PB/PCs to secrete high-affinity antibodies (Inoue et al., 2018). In addition, memory B cells can be recruited into newly formed GCs to further mature upon secondary antigen exposure (Inoue et al., 2018). Overall, these mechanisms work in concert to establish antibody-mediated immune protection against infections.

Since the outbreak of SARS-CoV-2, numerous studies on human antibody responses against spike protein of this emerging virus have been conducted to inform the development of vaccines and antibody therapeutics (Lapuente et al., 2024; Qi et al., 2022; Roltgen and Boyd, 2024). Early in the pandemic, extensive isolation and characterization of monoclonal antibodies (mAbs) from convalescents reveals that neutralizing antibodies primarily target epitopes on receptor-binding domain (RBD) and to a lesser extent on N-terminal domain (NTD) of spike (Ling et al., 2023; Liu and Wilson, 2022). Furthermore, RBD-specific neutralizing antibodies are categorized into seven major groups according to their binding modes and fine epitopes (Hastie et al., 2021), whereas NTD-specific neutralizing antibodies largely target a single supersite (Cerutti et al., 2021; McCallum et al., 2021). Although spike-specific antibodies from different convalescents are highly diverse, a series of public antibodies (i.e. antibodies generated by different individuals while sharing common genes and antigen recognition modes) have been widely reported, including antibodies encoded by IGHV3-53, IGHV3-66, IGHV1-58 and IGHV1-69 (Robbiani et al., 2020; Wang et al., 2022a; Yan et al., 2022; Yuan et al., 2020a). In addition, longitudinal profiling of memory B cells from peripheral blood of convalescents demonstrates that spike- or RBD-specific memory B cells display increased somatic hypermutation, binding affinity and neutralizing potency over time, reflecting long-lasting GC responses elicited by primary SARS-CoV-2 infection (Gaebler et al., 2021; Sakharkar et al., 2021; Sokal et al., 2021). Compared with SARS-CoV-2 infection, two doses of SARS-CoV-2 mRNA vaccines lead to equivalent plasma neutralizing activity and similar amount of RBD-specific memory B cells (Wang et al., 2021). Spike- or RBD-specific mAbs cloned from vaccinees and convalescents recognize common epitopes and share frequently-used antibody genes (Wang et al., 2021). Moreover, analysis of draining lymph node samples from vaccinees indicates that mRNA vaccines provoke persistent GCs, which last for at least six months and enable the maturation of spike-specific B cells (Kim et al., 2022; Turner et al., 2021).

Under the immune pressure established by infection and vaccination, SARS-CoV-2 variants, epresented by Alpha (B.1.1.7), Beta (B.1.351), Gamma (P.1), Delta (B.1.617.2) and Omicron (B.1.1.529), emerged successively in the first two years of the pandemic. Among them, Omicron exhibited most striking evasion to neutralizing antibodies induced by ancestral SARS-CoV-2 spike and therefore caused a new wave of infections worldwide (Viana et al., 2022). Omicron infection in individuals administrated with mRNA vaccines predominantly recalls cross-reactive memory B cells elicited by vaccination whereas rarely induces *de novo* Omicron-specific B cells responses, highlighting that ancestral SARS-CoV-2 imprints B cell responses to Omicron (Kaku et al., 2022; Quandt et al., 2022; Wang et al., 2022b; Weber et al., 2023). Consequently, plasma neutralizing activity against ancestral SARS-CoV-2 and Omicron are enhanced synchronously (Kaku et al., 2022; Quandt et al., 2022; Wang et al., 2022b; Weber et al., 2023). Moreover, B cell clones with more mutations and higher affinities towards Omicron progressively remodel spike-specific memory B cell repertoire, suggesting that cross-reactive memory B cells can undergo further affinity maturation in new GCs induced by Omicron breakthrough infection (Kaku et al., 2023; Sokal et al., 2023).

With the rapid emergence of Omicron-derived variants, such as BA.2.75, BA.5, BF.7, BQ.1, XBB.1.5, XBB.1.16, EG.5.1, BA.2.86 and JN.1, repeated infections with these variants have become common and updated vaccines have been applied accordingly in some countries (Huang et al., 2023). In individuals with prior exposures to wild-type (WT) spike, antibody responses boosted by subsequent exposures to emerging Omicron variants, either through infection or vaccination, display biased neutralizing activities to ancestral SARS-CoV-2 and earlier Omicron variants, suggesting that immune imprinting may impede the elicitation of neutralizing antibodies against more recent and future variants (Johnston et al., 2024; Liang et al., 2024; Tortorici et al., 2024; Wang et al., 2023a; Yisimayi et al., 2024). Nonetheless, mAbs from those individuals dramatically surpass mAbs elicited by WT spike in neutralizing breadth and potency, indicating that immune imprinting is potentially beneficial for the development of broadly neutralizing antibodies (bnAbs) (Paciello et al., 2024; Sokal et al., 2023).

To better understand the beneficial effects of ancestral SARS-CoV-2 immune imprinting on antibody responses to emerging variants, we ask that to what extent imprinted B cells (i.e. memory B cells elicited by ancestral SARS-CoV-2) can develop neutralizing breadth and potency in immune recalls. To address this question, we performed a 7-month longitudinal study of B cells recognizing WT spike in two individuals of mRNA vaccine, from convalescence of breakthrough infection to acute phase of reinfection. Functional and genetic analysis of 632 mAbs from those B cells reveals that mAbs cloned after reinfection have dramatically enhanced neutralizing breadth and potency, which is attributed to recall of pre-existing memory B cells and maturation of those cells either before or after reinfection. Among the 11 mAbs that potently neutralize all tested SARS-CoV-2 variants from WT to KP.3, 5 mAbs are classified into public clonotypes encoded by IGHV3-53 or IGHV3-66, whereas the rest belong to a rarely reported clonotype encoded by IGHV3-74. Notably, IGHV3-74 mAbs can also broadly neutralize a panel of sarbecoviruses including SARS-CoV-1, Pangolin CoV GD, Bat CoV WIV16 and RaTG13. Structural and functional analysis shows that IGHV3-74 mAbs can block the binding between RBD and ACE2 by targeting a novel epitope consisting of E340-R346, V367-S375, N436-G446 and Q498-V503. Moreover, the best IGHV3-74 mAb, termed KXD355, is highly resilient to variations on this epitope. Overall, this study demonstrates that both public and rare antibody clonotypes imprinted by ancestral SARS-CoV-2 have potential to achieve extraordinary neutralizing breadth and potency in repeated Omicron infections, which implicates that ancestral SARS-CoV-2 immune imprinting can be harnessed in developing pan-SARS-CoV-2 and even pan-sarbecovirus vaccines.

## Results

### Reinfection enhances plasma neutralizing antibody responses

In our previous study, we recruited a cohort to study antibody responses elicited by breakthrough infection (Li et al., 2023). Among them, Donor 1 and 2 received two doses of mRNA vaccines in the middle or beginning of 2021 and were infected by BA.5 or BF.7 in the end of 2022 (Figure 1A). With their blood samples collected about two months (T1) after infection, we have identified multiple mAbs with broadly neutralizing capacity (Li et al., 2023). To longitudinally track B cell responses in these two donors, we also collected blood samples about 3 months (T2) and 6 months (T3) after infection. Donor 1 showed symptoms of infection and was confirmed by SARS-CoV-2 antigen test on August 25, 2023, when EG.5.1 caused a new round of infections in China (Wang et al., 2023b). Eleven days later (T4), blood was collected again from Donor 1 to study acute phase antibody responses evoked by reinfection. In parallel, we also collected blood from Donor 2 at T4, though Donor 2 reported no symptoms and negative test results during several waves of infections before this time point.

**Figure 1.**
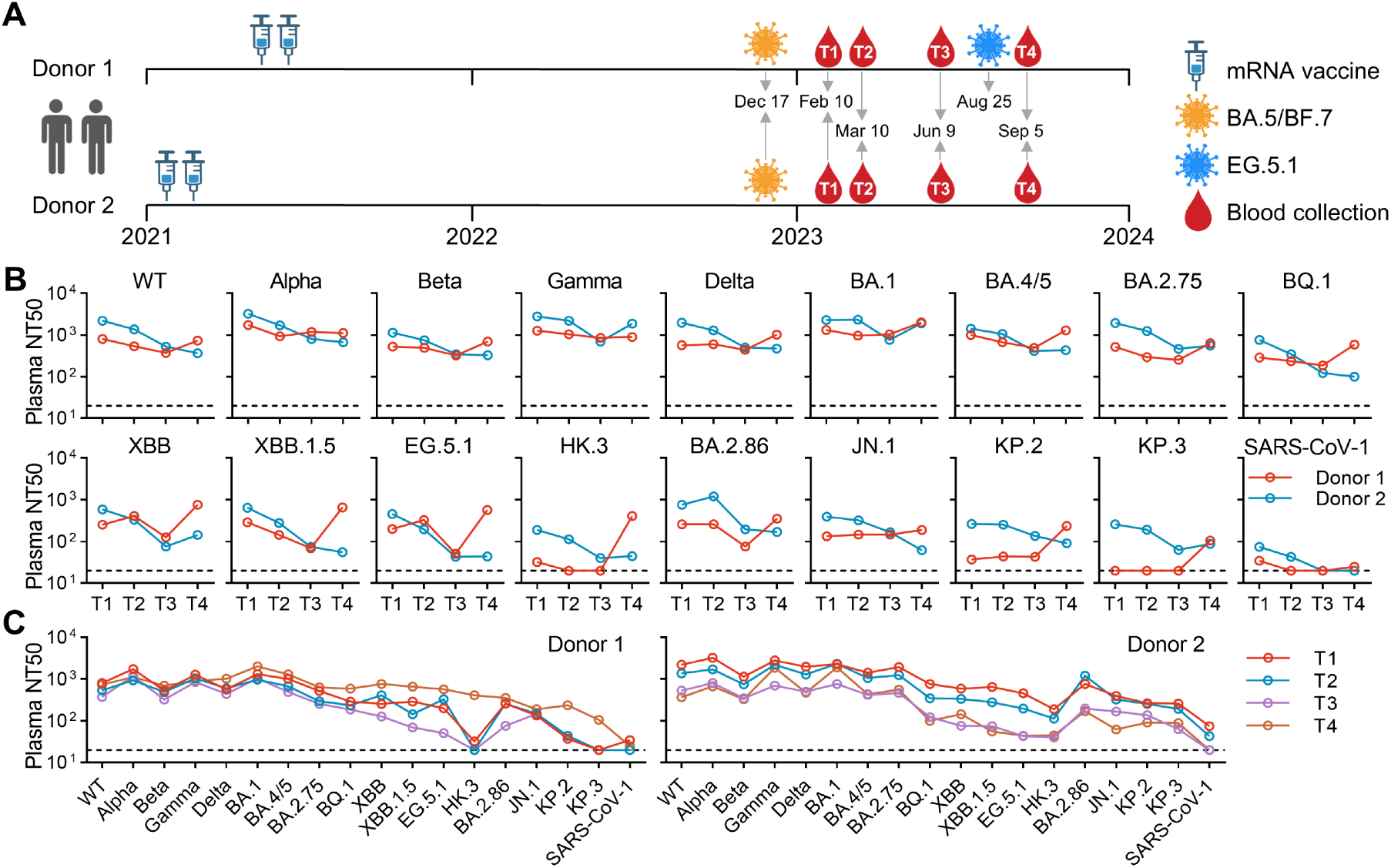
Longitudinal analysis of plasma neutralizing antibody responses in two individuals. (A) Basic information of donors and samples. (B, C) Summary of plasma neutralizing antibody titers (NT50). NT50s are calculated with results from three independent experiments, in which duplicates are performed.

We measured neutralizing antibody titers of plasma from four time points against a panel of 18 pseudoviruses bearing spikes from ancestral SARS-CoV-2 (WT), Alpha, Beta, Gamma, Delta, BA.1, BA.4/5, BA.2.75, BQ.1, XBB, XBB.1.5, EG.5.1, HK.3, BA.2.86, JN.1, KP.2, KP.3 and SARS-CoV-1 (Figure 1B and Figure S1). Overall, the plasma neutralizing titers (NT50s) of both donors gradually decrease from T1 to T3 with a maximum of 10.7 folds. After reinfection, plasma from Donor 1 at T4 shows substantially increased neutralizing titers against most strains by a maximum of 20.7 folds compared to T3. The titers of plasma from Donor 2 continuously decline or sustain from T3 to T4 for most strains, whereas increase by 1.9 to 2.7 folds for strains including Gamma, BA.1 and XBB. As latest variants of XBB and BA.2.86 lineages, HK.3 and KP.3 are least sensitive to neutralization, highlighting the continuous neutralization escape (Figure 1C). Nonetheless, plasma at T4 from Donor 1 displays relatively stable neutralizing titers across all SARS-CoV-2 variants, indicating that reinfection enhances the breadth of neutralizing antibody response in Donor 1.

### Monoclonal antibodies cloned from WT-spike-specific B cells after reinfection exhibit improved neutralizing breadth and potency

As B cells imprinted by ancestral SARS-CoV-2 mRNA vaccine (i.e. memory B cells) typically maintain their specificities to WT spike in recalled responses (Johnston et al., 2024; Kaku et al., 2022; Kaku et al., 2023; Liang et al., 2024; Quandt et al., 2022; Sokal et al., 2023; Tortorici et al., 2024; Wang et al., 2023a; Wang et al., 2022b; Weber et al., 2023; Yisimayi et al., 2024), we used WT spike as bait to track imprinted B cells in peripheral blood mononuclear cells (PBMCs) from these two donors. Consistent with plasma neutralizing titers, the frequencies of WT-spike-specific B cells in Donor 1 drop sequentially from T1 to T3 and then rise back to initial level (Figure 2A). Unexpectedly, Donor 2 displays similar dynamic changes as Donor 1, suggesting that Donor 2 may have experienced an asymptomatic infection between T3 and T4.

**Figure 2.**
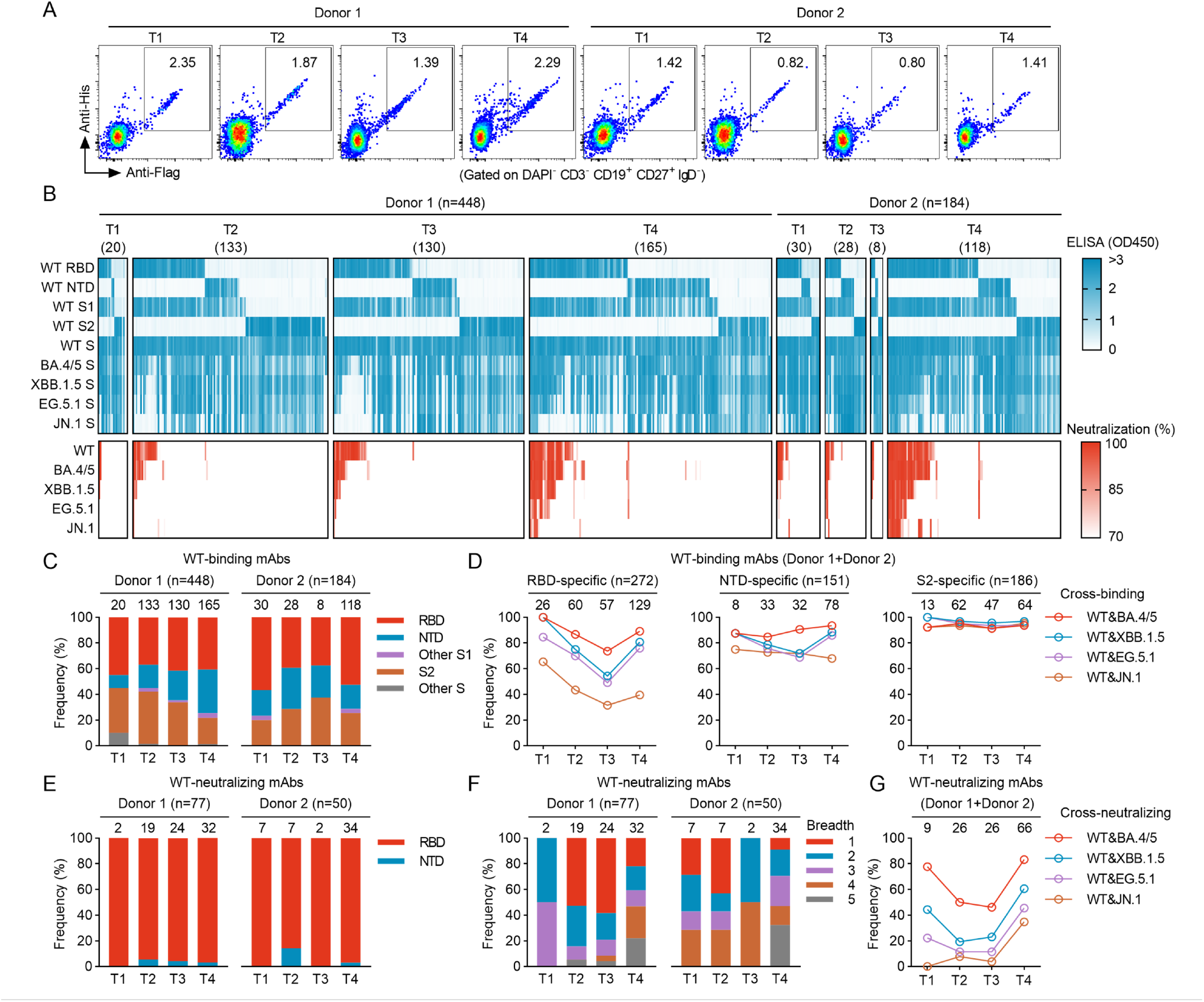
Isolation and characterization of mAbs. (A) FACS plots representing the percentages of WT-spike-specific B cells in DAPI^-^CD3^-^CD19^+^CD27^+^IgD^-^ B cells. (B) ELISA and neutralization results of 632 mAbs from two donors. Supernatant of HEK293T is tested against 9 antigens by ELISA, and tested against 5 pseudoviruses by neutralization assay. For ELISA, supernatant is not diluted. For neutralization, supernatant is diluted by 4.5 folds. The results are mean values from two independent experiments, in which duplicates are performed. (C) Frequencies of mAbs targeting each domain among mAbs binding to WT spike (WT-binding mAbs). (D) Frequencies of cross-binding mAbs among RBD-specific, NTD-specific, and S2-specific mAbs. (E) Frequencies of mAbs targeting RBD or NTD among mAbs neutralizing WT pseudoviruses (WT-neutralizing mAbs). (F) Frequencies of mAbs with different neutralizing breadth. Breadth indicates the number of pseudoviruses can be neutralized by mAbs. (G) Frequencies of cross-neutralizing mAbs.

To examine the binding and neutralizing properties of antibodies from those WT-spike-specific B cells, we cloned the variable regions of antibodies from those B cells into expression vectors through single-cell RT-PCR. Then we transfected HEK293T cells with those vectors and harvested supernatant for ELISA and neutralization assay (Figure 2B). In ELISA, the supernatant was tested with RBD, NTD, S1, S2 and spike (S) from WT SARS-CoV-2 and spikes from Omicron variants including BA.4/5, XBB.1.5, EG.5.1 and JN.1. In neutralization assay, the supernatant was tested against WT, BA.4/5, XBB.1.5, EG.5.1 and JN.1 pseudoviruses.

With published mAbs at T1 included, we identified 448 and 184 WT-spike-specific mAbs from Donor 1 and Donor 2 respectively. The main targets of these mAbs are RBD, NTD and S2, with the proportions of mAbs to each domain slightly varying from T1 to T4 (Figure 2C). Regarding to cross-binding with different spikes, the frequencies of RBD-specific mAbs that bind WT and one Omicron variant, either BA.4/5, XBB.1.5, EG.5.1 or JN.1, decline from T1 to T3 and then ascend (Figure 2D, Figure S2A). However, similar trend is not observed in NTD- and S2-specific mAbs.

By defining 70% inhibition as cutoff, we identified 127 mAbs that are able to neutralize WT pseudoviruses (WT-neutralizing mAbs), with 77 from Donor 1 and 50 from Donor 2 (Figure 2E). Among them, only 5 mAbs target NTD while the others exclusively recognize RBD. Regarding to cross-neutralizing activity, around 20-30% of mAbs at T4 are able to neutralize all 5 tested pseudoviruses, whereas only 2 out of 61 mAbs from T1 to T3 have such capacity (Figure 2F). Moreover, the frequencies of mAbs that cross-neutralize WT and one Omicron variant drop from T1 to T3 and then jump to highest levels at T4 (Figure 2G, Figure S2B).

To further compare the breadth and potency of neutralizing mAbs from different time points, we purified those mAbs and measured their IC50s against 14 representative pseudoviruses including WT, Delta, BA.1, BA.4/5, BA.2.75, BQ.1, XBB.1.5, EG.5.1, HK.3, BA.2.86, JN.1, KP.2, KP.3 and SARS-CoV-1 (Figure 3A). Four neutralizing mAbs screened by supernatant were not included in this analysis because of low yields. Among all tested mAbs, 4 mAbs from Donor 2 are able to neutralize all SARS-CoV-2 variants and SARS-CoV-1. In addition, 2 mAbs from Donor 1 and 5 mAbs from Donor 2 can neutralize all SARS-CoV-2 variants from WT to KP.3. Notably, these 11 mAbs are exclusively identified from PBMCs collected at T4. Along with this finding, differences are observed in the overall neutralizing breadth of mAbs at T1 to T4 (Figure 3B, Figure S3A). T1 and T4 have higher frequencies of neutralizing antibody against BA.1, BA.4/5, BA.2.75, BQ.1, XBB.1.5 and BA.2.86 than T2 and T3, which is consistent with neutralization results of supernatant. Moreover, T4 has higher proportions of neutralizing antibody against EG.5.1, HK.3, JN.1, KP.2 and KP.3 than the other time points.

**Figure 3.**
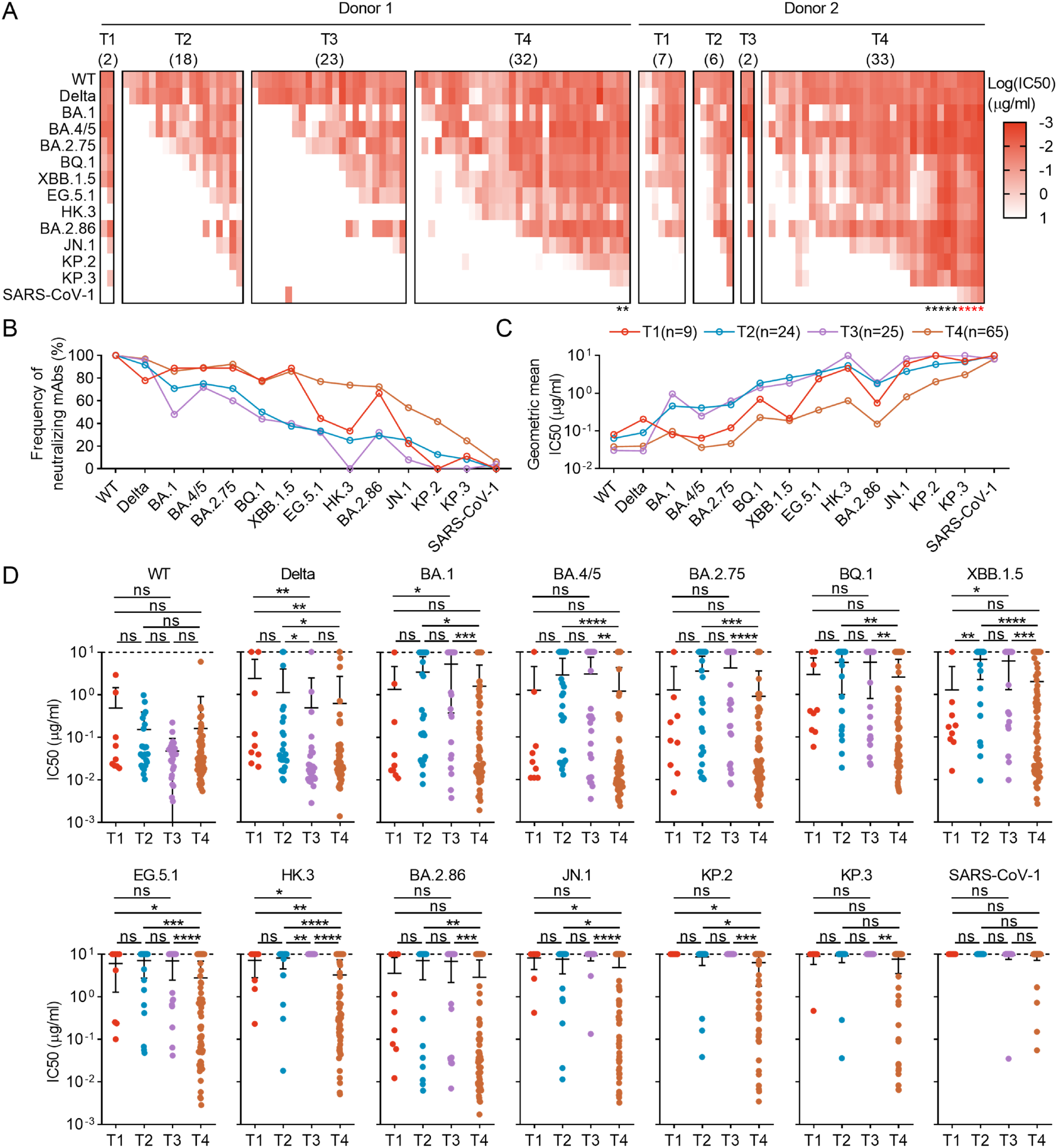
Neutralizing potency of mAbs against SARS-CoV-2 and SARS-CoV-1. (A) IC50s of mAbs against 14 pseudoviruses. The highest antibody concentration used to determine IC50 is 10 μg/ml. IC50s are calculated with results from two independent experiments, in which duplicates are performed. The black asterisk indicates mAbs with broadly neutralizing activity against SARS-CoV-2 variants from WT to KP.3. The red asterisk indicates mAbs with broadly neutralizing activity against SARS-CoV-2 and SARS-CoV-1. (B) Frequencies of neutralizing mAbs against each pseudoviruses among mAbs from T1 to T4. (C) Geometric mean IC50s of mAbs from T1 to T4 against each pseudoviruses. (D) Comparison of IC50s. Statistical analysis is performed by Mann-Whitney test. ns (not significant) P > 0.05, *P < 0.05, **P < 0.01, ***P < 0.001, ****P < 0.0001.

Regarding to neutralizing potency, the overall IC50s of mAbs at T4 are lower than that of mAbs at T2 and T3 against most strains (Figure 3C, D, Figure S3A). In addition, mAbs at T4 also display superior neutralizing potency than mAbs at T1 against Delta, EG.5.1, HK.3, JN.1 and KP.2. In contrast, significant differences in IC50s are generally not observed among mAbs at T1 to T3 by statistical analysis. We also combined mAbs at T1 to T3 together and compared them with mAbs at T4, which further confirms that mAbs isolated after reinfection have superior neutralizing breadth and potency than mAbs before reinfection (Figure S3B-D). Taken together, these data suggest that mAbs cloned from WT-spike-specific B cells after reinfection have enhanced neutralizing breadth and potency.

### Reinfection recalls pre-existing memory B cells

To explain the enhanced neutralizing breadth and potency of mAbs cloned after reinfection, we moved on to analyze the sequences of mAbs from T1 to T4. Binding mAbs from both donors use a wide range of V genes (Figure 4A, B). The frequencies of each V gene from T1 to T4 are more consistent for mAbs from Donor 1 than those from Donor 2, likely due to fewer mAbs isolated from Donor 2 at T2 and T3. The most frequently-used V genes for heavy chain include IGHV3-30, IGHV3-66, IGHV3-53, IGHV3-33, IGHV1-69 and IGHV1-46. Among them, IGHV3-30 and IGHV3-33 are primarily used by mAbs targeting S2, whereas IGHV3-66, IGHV3-53 and IGHV1-69 are preferentially used by mAbs recognizing RBD (Figure S4A). The most frequently-used V genes for light chain include IGKV1-39, IGKV3-20, IGKV3-15, IGKV3-11, IGKV1-33, IGKV1-5 and IGKV1-9, which are less biased toward mAbs targeting a certain domain than V genes for heavy chain (Figure S4A). In contrast to binding mAbs, neutralizing mAbs are predominantly encoded by IGHV3-66, IGHV3-53 and IGHV1-69 (Figure 4A, B). Furthermore, in these neutralizing mAbs, IGHV3-66 and IGHV3-53 typically pair with IGKV1-33, IGKV1-9 and IGKV3-20, whereas IGHV1-69 commonly pair with IGLV1-40 (Figure S4B). It is worth mentioning that the pair of IGHV3-74 and IGKV3-20 is used by 10 out of 50 neutralizing mAbs from Donor 2. Nonetheless, the overall profiles of V gene usage are rather consistent from T1 to T4. According to V and J genes and CDR3 similarity, we defined 401 and 145 unique clonotypes respectively from 448 and 184 mAbs (Figure 4C). The expanded clonotypes are primarily found in mAbs at T4, particular for Donor 2, further indicating that T4 is also in acute phase of reinfection for Donor 2. As an evidence of recall of pre-existing memory B cells, multiple clonotypes found at T1 to T3 are extensively expanded at T4, particular for neutralizing clonotypes such as Clonotype 204 and 223 from Donor 1 and Clonotype 88 and 92 from Donor 2 (Figure 4D).

**Figure 4.**
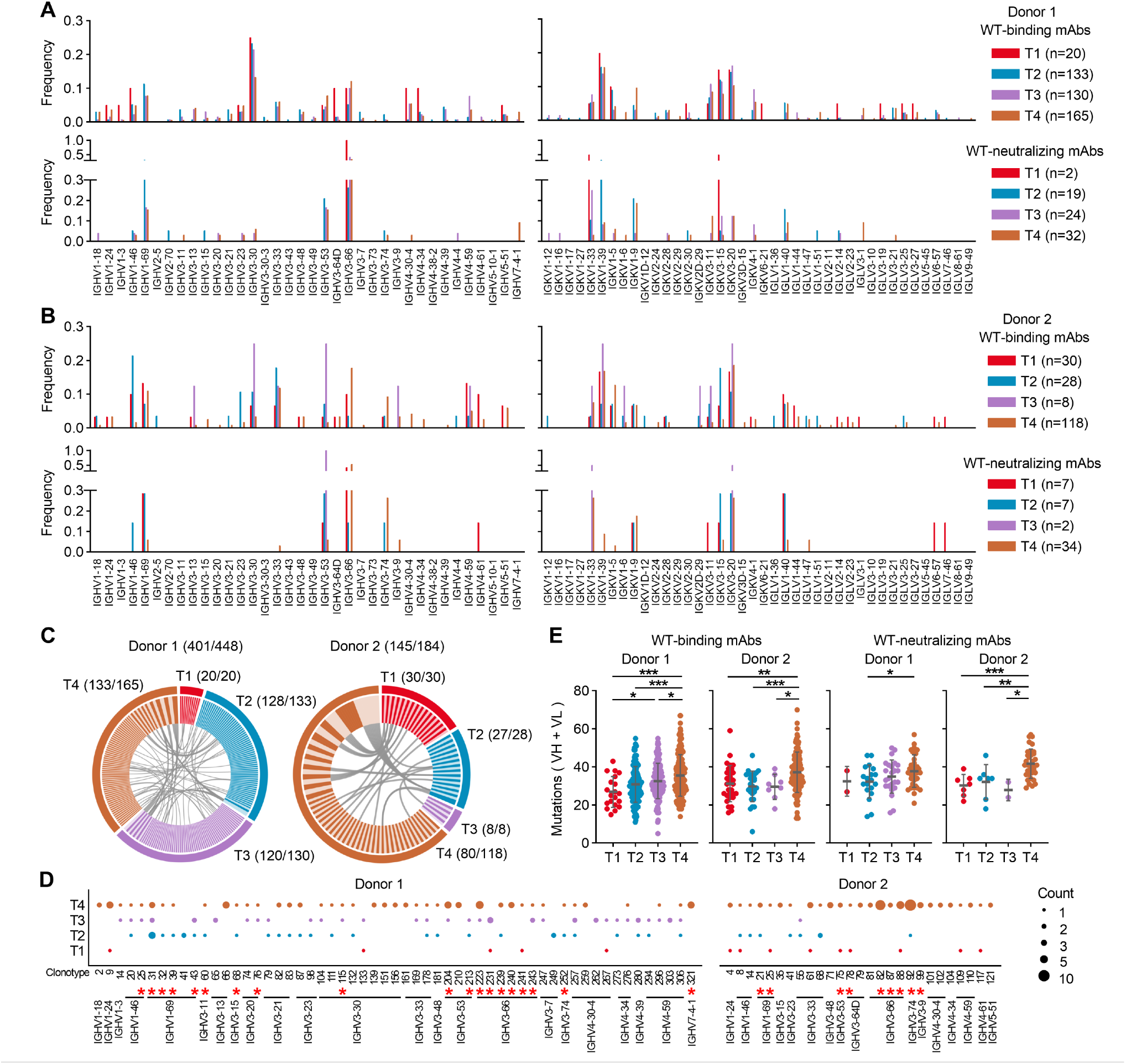
Sequence analysis. (A, B) Frequencies of V genes used by mAbs binding to WT spike (WT-binding mAbs) and mAbs neutralizing WT pseudoviruses (WT-neutralizing mAbs) from Donor 1 and Donor 2. (C) Clonotype analysis of WT-binding mAbs from Donor 1 and Donor 2. Values in parentheses indicate number of clonotypes/number of mAbs. Different clonotypes are distinguished by alternating colors, and identical clonotypes across different time points are connected by gray lines. (D) Bubble plot of clonotypes comprising two or more mAbs. Dot size indicates number of mAbs. The red asterisk indicates clonotypes containing neutralizing mAbs. (E) Somatic hypermutation on variable regions of heavy and light chains based on DNA sequences. Statistical analysis is performed by two-tailed unpaired T test. *P<0.05, **P < 0.01, ***P < 0.001.

On the other hand, mAbs at T4 have more somatic mutations than those from earlier time points, which can be either due to recall of pre-existing memory B cells with more mutations, or due to further maturation of pre-existing memory B cells in new GC reactions (Figure 4E). Taken together, these data suggest that the enhanced neutralizing breadth and potency of mAbs at T4 is attributed to recall of pre-existing memory B cells and maturation of those cells either before or after reinfection.

### Both public and rare antibody clonotypes can develop extraordinary neutralizing breadth and potency

After longitudinal analysis of overall B cell responses in these two donors, we moved on to characterize the 11 mAbs with extraordinary neutralizing breadth (Figure 3A, Figure 5A). Clonotype analysis reveals that they belong to 5 unique clonotypes including Clonotype 92, 82 and 75 from Donor 2 and Clonotype 240-2 and 212-1 from Donor 1. Clonotype 92 is encoded by IGHV3-74 and IGHV3-20, which is rarely reported in RBD-specific antibodies against SARS-CoV-2 (Wang et al., 2022a). This clonotype contains 10 neutralizing mAbs, with 1 from T2 and 9 from T4. Among them, KXD350, KXD352, KXD355 and KXD358 can neutralize all tested SARS-CoV-2 variants and SARS-CoV-1. Further neutralization analysis shows that KXD350 and KXD355 can also neutralize other sarbecoviruses including Pangolin CoV GD, Bat CoV WIV16 and RaTG13-T372A, whereas KXD352 and KXD358 are escaped by RaTG13-T372A (Figure 5B). According to the phylogenetic tree, mAbs in this clonotype can be divided into two major clades, suggesting that neutralizing breadth can be developed through diverse evolution pathways.

**Figure 5.**
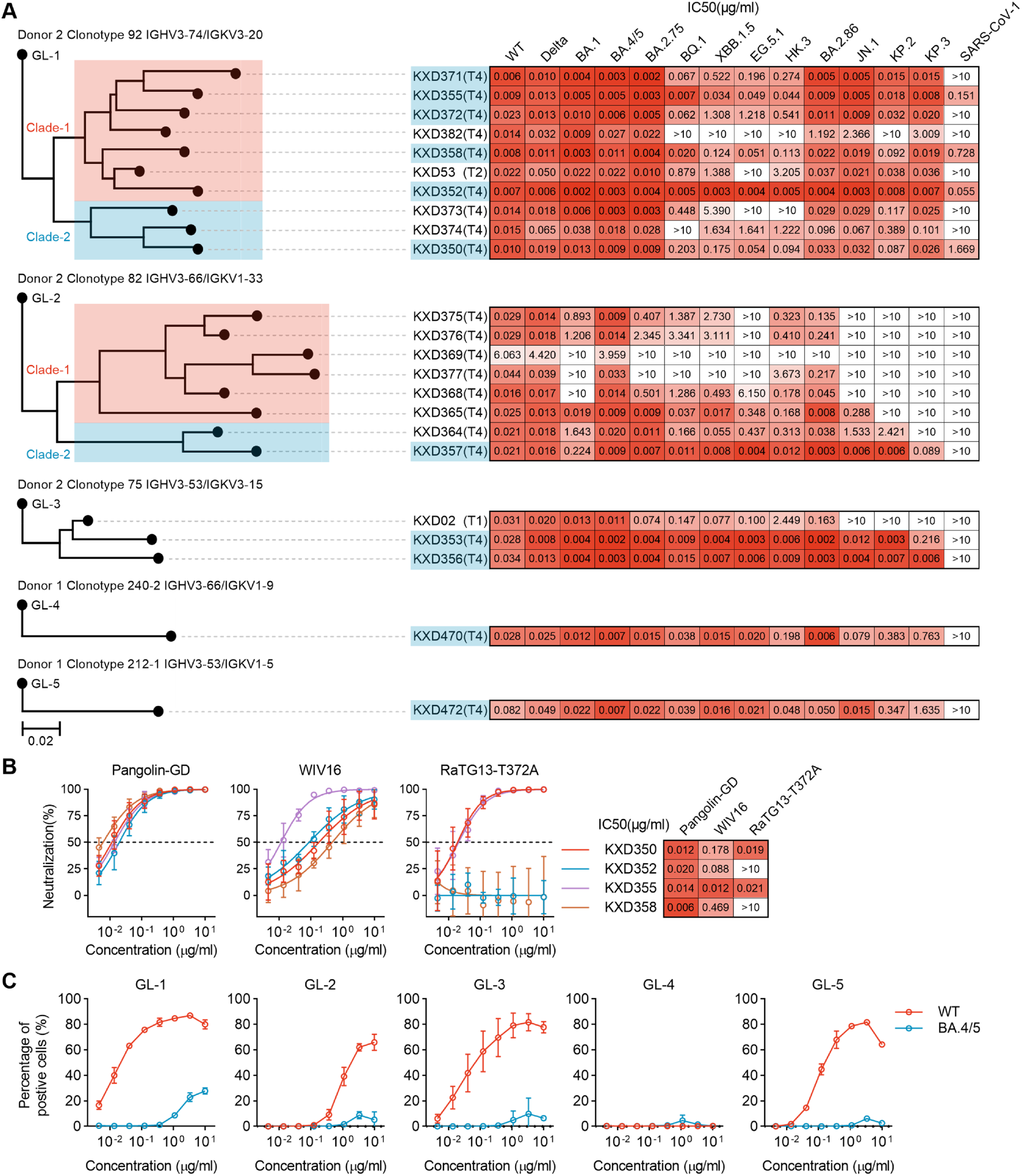
Clonotypes containing mAbs with extraordinary neutralizing breadth and potency. (A) Maximum-likelihood phylogenetic trees of each clonotype and IC50s of each mAbs. (B) Neutralization against sarbecoviruses including Pangolin CoV GD (Pangolin-GD), Bat CoV WIV16 (WIV16) and Bat CoV RaTG13-T372A (RaTG13-T372A). (C) Binding of germline antibodies to WT and BA.4/5 spikes expressed on HEK293T cells.

The other four clonotypes belong to previously defined IGHV3-53/3-66 public antibody clonotypes (Wang et al., 2022a). Clonotype 82 is encoded by IGVH3-66 and IGKV1-33 and contains 8 neutralizing mAbs from T4. However, only KXD357 is able to neutralize all SARS-CoV-2 variants, with IC50s ranging from 0.003 to 0.224 μg/ml. Similar as Clonotype 92, Clonotype 82 also has two major clades. Although mAbs in both clades are extensively mutated, mAbs in Clade-2 exhibit superior neutralizing breadth than mAbs in Clade-1. Encoded by IGHV3-53 and IGKV3-15, Clonotype 75 contains previously published KXD02 from T1 and two addition mAbs from T4. In our previous study, we have shown that KXD02 broadly neutralizes SARS-CoV-2 variants from WT to HK.3 (Li et al., 2023). However, KXD02 is soon escaped by variants in BA.2.86 lineages including JN.1, KP.2 and KP.3. In contrast, KXD353 and KXD356 from T4 maintain neutralizing activities against all SARS-CoV-2 variants, with IC50s ranging from 0.002 to 0.216 μg/ml. In line with increased neutralizing breadth and potency, KXD353 and KXD356 are also more evolved than KXD02 according to phylogenetic analysis. As singletons from Donor 1, Clonotype 240-2 (KXD470) is encoded by IGHV3-66 and IGKV1-9, whereas Clonotype 212-1 (KXD472) is encoded by IGHV3-53 and IGKV1-5. Although both mAbs are able to neutralize all SARS-CoV-2 variants, they are less potent to KP.2 and KP.3 than bnAbs from Donor 2.

As all these clonotypes are identified with WT spike, we assume that they are initially primed by ancestral SARS-CoV-2 mRNA vaccine but not by BA.5 or BF.7 breakthrough infection. To confirm this assumption, we synthesized the putative germline antibodies of each clonotypes and measured their binding avidities to WT and BA.4/5 spikes expressed on HEK293T cells (Figure 5C). BF.7 was not tested since it has only one amino acid different from BA.5 in RBD. All the germlines, except GL-4, show strong to moderate binding to WT spike. In contrast, they all display weak or even undetectable binding to BA.4/5 spike. The biased recognition towards WT spike suggest that these clonotypes are more likely to be activated by mRNA vaccine. Taken together, these data suggest that both public and rare antibody clonotypes imprinted by ancestral SARS-CoV-2 have potential to develop ultra-broad and potent neutralizing activities in repeated Omicron infections.

### IGHV3-74-encoded bnAbs target a novel epitope and exhibit dramatic resilience to mutations on the epitope

To evaluate the neutralizing potential of above bnAbs to future SARS-CoV-2 variants, we moved on to analyze their epitopes. We first performed competition ELISA to roughly map their epitopes with ACE2, CR3022, S309, P2S-2E9, BD55-5483, BD56-1854 and BD55-1205 as references (Figure 6A).

**Figure 6.**
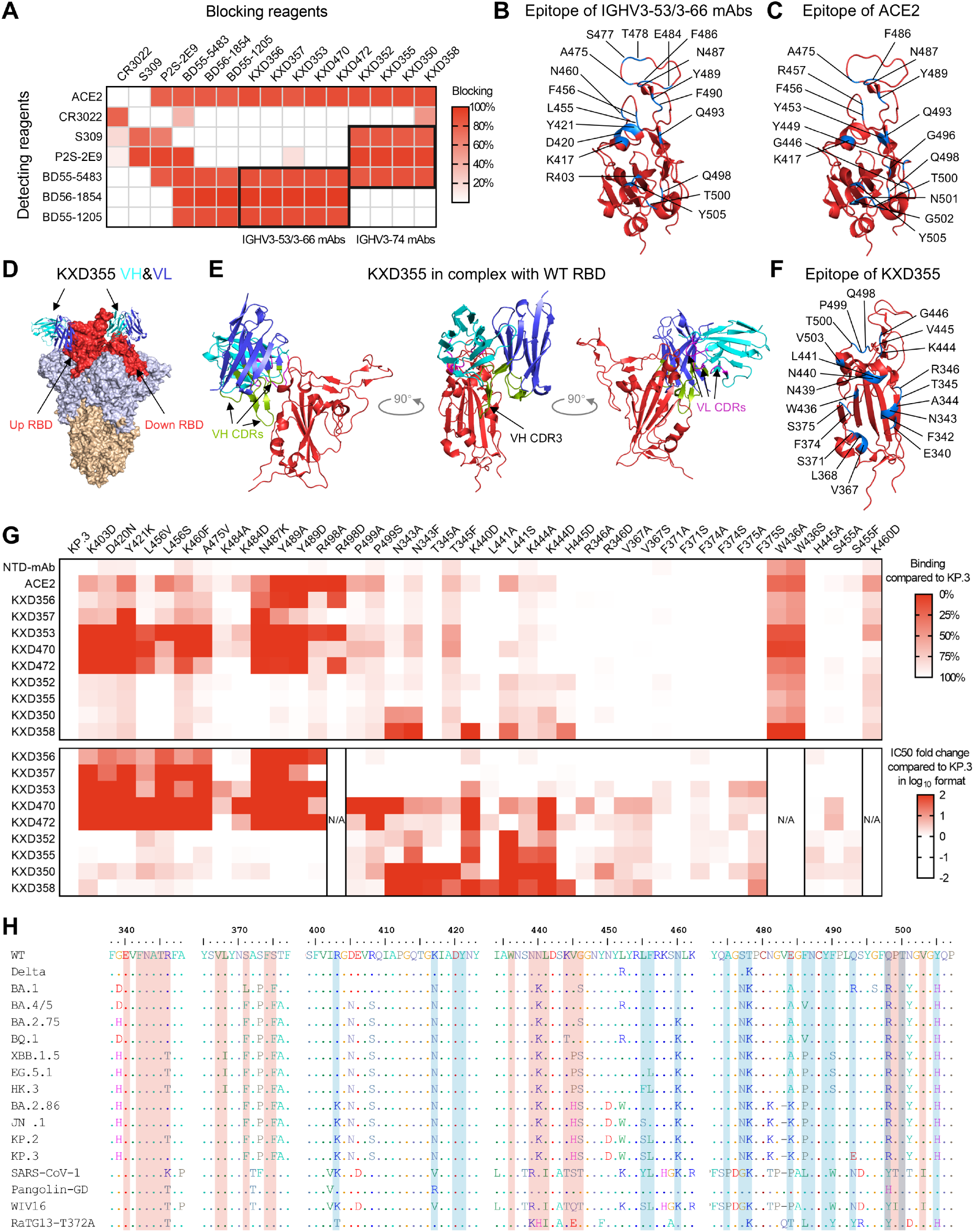
Epitope mapping and escape mutations of mAbs with extraordinary neutralizing breadth and potency. (A) Competition ELISA. The blocking indicates signal (OD450) reduced by blocking mAbs. The results are mean values from two independent experiments, in which duplicates are performed. (B) Epitope of IGHV3-53/3-66 mAbs on WT RBD (PDB: 7A94). (C) Epitope of ACE2 on WT RBD (PDB: 7A94). (D) The predicted structure of KXD355 in complex with WT RBD is aligned to up RBD and down RBD on WT spike (PDB: 7A94). (E) Detailed view of KXD355 in complex with WT RBD. (F) Epitope of KXD355 on WT RBD (PDB: 7A94). (G) Binding and neutralizing activities against single-mutated KP.3. The binding activities are measured by FACS. NTD-mAb, a mAb targeting NTD, is used to evaluate the expression levels of KP.3 mutants. The results of binding activities are mean values from two independent experiments. For neutralizing activities, IC50s are calculated with results from two independent experiments, in which duplicates are performed. N/A (not applicable) indicates that neutralization assay is not done because we are not able to generate pseudoviruses for the mutant. (H) Alignment of spike sequences from SARS-CoV-2 variants and other sarbecoviruses tested in neutralization assay. The epitopes of IGHV3-53/3-66 and IGHV3-74 mAbs are respectively highlighted with blue and red background.

As reported, CR3022 binds to a cryptic epitope on the inner face of RBD whereas S309 recognizes N343 proteoglycan epitope on the outer face of RBD (Figure S5A) (Pinto et al., 2020; Yuan et al., 2020b). P2S-2E9 and BD55-5483 (analogue of BD55-5514) both target the previously defined “mesa” of RBD, respectively biased to the outer face and inner face (Cao et al., 2022; Hastie et al., 2021; Ju et al., 2023). As representatives of IGHV3-53/3-66 public antibodies, BD56-1854 and BD55-1205 target receptor-binding-motif (RBM) (Cao et al., 2023).

We tested four mAbs (KXD350, KXD352, KXD355 and KXD358) from Clonotype 92 (IGHV3-74 mAbs) and five mAbs (KXD353, KXD356, KXD357, KXD470 and KXD472) from other clonotypes (IGHV3-53/3-66 mAbs) (Figure 6A). IGHV3-53/3-66 mAbs compete with BD55-5483, BD56-1854 and BD55-1205 but not with S309 and P2S-2E9. In contrast, IGHV3-74 mAbs compete with S309, P2S-2E9 and BD55-5483 while not with BD56-1854 and BD55-1205. Moreover, no competition is observed between these mAbs and CR3022 except for some minor competition between KXD358 and CR3022. In addition, they all compete with ACE2, suggesting that they neutralize SARS-CoV-2 through receptor blockade.

To define the fine epitopes of these mAbs, we performed structure prediction with AlphaFold 3 (Abramson et al., 2024). The predicted structures of IGHV3-53/3-66 mAbs in complex with WT RBD are similar to the structure of BD55-1205 in complex with XBB.1.5 RBD (Figure S5B). According to these structures, we defined the common epitope of IGHV3-53/3-66 mAbs, which largely overlap with the epitope of ACE2 (Figure 6B, C).

On the other hand, AlphaFold 3 generated various structures for the complexes of IGHV3-74 mAbs and WT RBD. However, only one type of structures is consistent with epitope mapping by competition ELISA. Here, KXD355 is taken as an example to present the results. Binding to the previously defined “escarpment” of RBD (Hastie et al., 2021), KXD355 clashes with S309, P2S-2E9 and BD55-5514 (analogue of BD55-5483) but not with BD55-1205 and CR3022 (Figure S5C). As this epitope is exposed on both up and down RBDs, KXD355 can bind RBDs in both conformations (Figure 6D). In details, the long CDRH3 of KXD355 extends into a groove formed by three helixes on the “escarpment” of RBD (Figure 6E). Meanwhile, the other CDRs attach to “mesa” of RBD to provide support for CDRH3. Accordingly, the epitope of KXD355 includes 22 residues distributed on E340-R346, V367-S375, W436-G446 and Q498-V503 (Figure 6F).

To confirm the predicted epitopes, we constructed a panel of single-mutated KP.3 spikes and measured their escape to those mAbs in binding and neutralization (Figure 6G). Compared to KP.3, most mutants show similar or slightly lower expression levels on HEK293T cells, which is indicated by an NTD-specific mAb. In contrast, nearly half of these mutants display reduced binding to ACE by different extent. Regarding to the binding with mAbs, mutants including K403D, D420N, Y421K, L456V, L456S, K460F, A475V, N487K, Y489A, Y489D, R498A and R498D completely lose binding to at least one IGHV3-53/3-66 mAb. To a lesser extent, mutants like N343A, N343F, K440D, L441S and H445D show decreased binding to IGHV3-74 mAbs. Consistent with reduced binding, the aforementioned mutants also display dramatic escape of neutralization to corresponding mAbs. In addition, mutants such as T345A, T345F, K444A and K444D are also resistant to certain IGHV3-74 mAbs, although they only partially lose binding to those mAbs.

In contrast to all IGHV3-53/3-66 mAbs and other IGHV3-74 mAbs, KXD352 and KXD355 maintain low IC50s against all the mutants except L441S, suggesting that they are more resilient to single mutations on their epitopes (Figure S6). Compared with SARS-CoV-2 variants, SARS-CoV-1 and other sarbecoviruses have multiple mutations on the epitopes of IGHV3-53/3-66 and IGHV3-74 mAbs (Figure 6H). However, the best IGHV3-74 mAb, KXD355 can neutralize all the tested sarbecoviruses with IC50s ranging from 0.003 to 0.151 μg/ml, further highlighting that KXD355 is highly tolerant to variations on its epitope (Figure 5A, B).

To evaluate the likelihood that these bnAbs are escaped by future circulating SARS-CoV-2 strains, we collected 25332 spike sequences published between June 1 to October 1 in 2024. With KP.3 as reference, we analyzed the mutations on the epitopes of these bnAbs. With extremely low frequency, multiple confirmed escape mutations for IGHV3-53/3-66 mAbs are found in the circulating strains, including D420N (0.008%), A475V (0.288%), N487K (0.008%) (Figure S7A). In contrast, confirmed escape mutations for IGHV3-74 mAbs are not found in the circulating strains (Figure S7B). Overall, these data predict that IGHV3-74 mAbs, particular KXD355, will not be easily escaped by SARS-CoV-2 in near future.

## Discussion

The key scientific question of this study is to what extent B cells imprinted by ancestral SARS-CoV-2 can develop neutralizing breadth and potency in immune recalls. Strategically, we used WT spike as bait to track those B cells as they typically maintain specificities to WT spike in immune recalls (Johnston et al., 2024; Kaku et al., 2022; Kaku et al., 2023; Liang et al., 2024; Quandt et al., 2022; Sokal et al., 2023; Tortorici et al., 2024; Wang et al., 2023a; Wang et al., 2022b; Weber et al., 2023; Yisimayi et al., 2024). A recent study pointed out that memory B cells induced by ancestral SARS-CoV-2 could redirect their specificities to emerging variants in repeated Omicron exposures (Kotaki et al., 2024). As our focus was memory B cells broadly recognizing SARS-CoV-2 from WT to emerging variants, we did not take into account those memory B cells with redirected specificities. Regarding to the samples, we collected blood from 2 individuals of mRNA vaccines at four time points, which are 2 months, 3 months, 6 months and 9 months after BA.5 or BF.7 breakthrough infection. In addition, Donor 1 experienced a symptomatic reinfection at 11 days before the last time point, while Donor 2 experienced an asymptomatic reinfection shortly before the last time point, as suggested by our data. Therefore, the timeline of this longitudinal study is ranging from convalescence of breakthrough infection to acute phase of reinfection.

A major finding of this study is that mAbs cloned after reinfection have superior neutralizing breadth and potency, which is accompanied with expansion of pre-existing clonotypes and increased mutation levels. According to the antigen exposure history of these two donors, this finding can be explained by two possibilities. First, a fraction of pre-existing memory B cells, which have already undergone extensive maturation and developed superior neutralizing breadth and potency, are recalled by reinfection. Second, a fraction of pre-existing memory B cells are recalled by reinfection and then acquire further maturation in new GC responses. Non-mutually exclusive, these two possibilities differ in whether an addition round of maturation occurs after reinfection. Previous studies suggest that BA.1 breakthrough infection or booster vaccination can both lead to affinity maturation of pre-existing memory B cells induced by mRNA vaccines (Alsoussi et al., 2023; Kaku et al., 2023; Sokal et al., 2023). Here, it remains to be determined whether memory B cells induced by mRNA vaccines can undergo two rounds of affinity maturation sequentially driven by breakthrough infection and reinfection.

A highlight of this study is the identification of 4 bnAbs against sarbecoviruses. These bnAbs belong to a clonotype encoded by IGHV3-74, which is rarely used by RBD-specific antibodies (Wang et al., 2022a). Structural analysis indicates that these IGHV3-74 bnAbs bind to an epitope consisting of E340-R346, V367-S375, N436-G446 and Q498-V503. In an early study, epitopes on RBD are classified into seven groups, with RBD-1 to RBD-3 on the top face, RBD-4 and RBD-5 on the outer face, RBD-6 and RBD-7 on the inner face (Hastie et al., 2021). More recently, a cryptic epitope recognized by human mAb BIOLS56 and IMCAS74 is defined as RBD-8 (Rao et al., 2023). Referred to these epitope categories, the epitope of these IGHV3-74 bnAbs stretches across RBD-3 and RBD-5 and is therefore a novel epitope. Unexpectedly, this epitope is not conserved among sarbecoviruses although IGHV3-74 bnAbs can broadly neutralize those viruses. According to structures predicted by AlphaFold 3, we speculate that the resilience of IGHV3-74 bnAbs to epitope variations is due to the plasticity of their long CDRH3s, which play a key role in contacting with this epitope.

As a byproduct of defining a novel broadly neutralizing epitope on RBD, our study represents an example of epitope mapping with AlphaFold 3. In our practice with AlphaFold 3, it is straightforward to predict the structures of IGHV3-53/3-66 mAbs, probably because the structures of similar antibodies have been abundantly reported. In contrast, various structures are predicted for IGHV3-74 bnAbs. In this case, traditional epitope mapping through competition ELISA is necessary to pick the best candidates for further confirmation. Overall, our attempt to apply AlphaFold 3 in fine epitope mapping turned out to be successful.

Similar to a study published recently (Jian et al., 2024), we identified multiple IGHV3-53/3-66 public antibodies with ultra-broad neutralizing activities against SARS-CoV-2. In addition, we observed the expansion of clonotypes encoded by IGHV3-53/3-66. According to phylogenetic analysis, the improved neutralizing breadth and potency of IGHV3-53/3-66 public antibodies is somehow not always correlated to the extent of somatic hypermutation, supporting the view that bystander mutations generated in a stochastic manner are critical for the development of bnAbs against SARS-CoV-2 (Bruhn et al., 2024; Korenkov et al., 2023). On the other hand, the neutralizing breadth of IGHV3-53/3-66 public antibodies reported so far is still limited to SARS-CoV-2 variants. It remains to be investigated whether this class of public antibodies can develop pan-sarbecovirus neutralization after further maturation.

Currently, ancestral SARS-CoV-2 immune imprinting has been acknowledged as a great challenge for the update of SARS-CoV-2 vaccines (Johnston et al., 2024; Liang et al., 2024; Tortorici et al., 2024; Yisimayi et al., 2024). In one preprint we posted recently, we find that immune imprinting can facilitate the development of bnAbs against SARS-CoV-2 (Wang et al., 2024 Preprint). Here, we further show that B cells imprinted by ancestral SARS-CoV-2 can even develop pan-sarbecovirus neutralization. Taken together, these studies support that ancestral SARS-CoV-2 immune imprinting can be harnessed in developing pan-SARS-CoV-2 and even pan-sarbecovirus vaccines. From the aspect of antibody therapeutics, KXD355 exhibits broad and potent neutralizing activity against sarbecoviruses. More importantly, KXD355 targets a novel epitope and tolerates mutations on this epitope. Therefore, KXD355 is an ideal candidate for antibody therapeutics against future SARS-CoV-2 variants and related sarbecoviruses.

## Materials and methods

### Ethics statement

This study was approved by the Ethics Committee of Hefei Institutes of Physical Science, Chinese Academy of Sciences (Approval Number: YXLL-2023-47). All donors provided written informed consent for collection of information, analysis of plasma and PBMCs, and publication of data generated from their samples.

### Human samples

Peripheral blood samples were collected from 2 donors. Plasma and PBMCs were isolated from blood through Ficoll density gradient centrifugation.

### Protein expression and purification

The extracellular domains of spikes from WT, BA.4/5, XBB.1.5, EG.5.1, JN.1 were constructed with a foldon-trimerization-motif, a His-tag, a tandem Strep-tag II and a FLAG-tag at C-terminal. S1, NTD and RBD of WT spike were constructed with a Strep-tag II at C-terminal. S2 of WT spike was constructed with a His-tag at C-terminal. Human ACE2 (huACE2, 1-740) was constructed with a Strep-tag II at N-terminal. All recombinant proteins were expressed in FreeStyle 293F cells and purified with Streptactin Agarose Resin 4FF (Yeasen, 20495ES60) or Ni-NTA Agarose (Qiagen, 30210).

### Cloning and expression of mAbs

As previously described (Li et al., 2023), PBMCs were first incubated with 200nM WT spike. After wash, cells were stained with antibodies targeting CD3, CD19, CD27, CD38, human IgM, human IgD, His-tag and FLAG-tag along with DAPI. WT-spike-specific B cells were gated as DAPI^-^ CD3^−^CD19^+^CD27^+^IgD^-^His^+^FLAG^+^ and sorted into 96-well PCR plates with one cell per well. After reverse transcription reaction, variable regions of antibody heavy and light chains from single B cells were amplified by nested PCR and cloned into human IgG1 expression vectors. Paired heavy and light chain plasmids were co-transfected into HEK293T cells or FreeStyle 293F cells. Antibodies were purified with Protein A magnetic beads (GenScript, L00273).

### ELISA

Indicated antigens were coated onto 96-well ELISA plates (100 ng/well) at 4°C overnight. After blocking with blocking buffer (PBS with 10% FBS), undiluted HEK293T supernatant was added to the wells and incubated at 37°C for 1hr. After wash, HRP-conjugated goat anti-human-IgG antibodies (Zen-bio, 550004) were added and incubated at 37°C for 1hr. After wash, TMB substrate (Sangon Biotech, E661007-0100) was added and incubated at room temperature for 5 minutes. Then the reaction was stopped with stop solution and absorbance at 450 nm (OD450) was measured.

### Pseudovirus neutralization assay

To generate pseudoviruses, HEK293T cells were transfected with psPAX2, pLenti-luciferase and spike-encoding plasmids (WT, Alpha, Beta, Gamma, Delta, BA.1, BA.4/5, BA.2.75, BQ.1, XBB, XBB.1.5, EG.5.1, HK.3, BA.2.86, JN.1, KP.2, KP.3, SARS-CoV-1, Pangolin CoV GD, Bat CoV WIV16, Bat CoV RaTG13 and KP.3 mutants) using polyetherimide (PEI). Pseudoviruses supernatants were collected 48hrs after transfection. 3-fold serially diluted plasma (starting at 1:20), HEK293T supernatant (starting at 1:4.5), or mAbs (starting at 10 μg/ml) were mixed with pseudoviruses at 37°C for 1hr. HEK293T-hACE2 cells (1.5×10^4^ per well) were added into the mixture and incubated at 37°C for 48hrs. Cells were lysed to measure luciferase activity by Bright-Lite Luciferase Assay System (Vazyme Biotech, DD1204-02). The percentages of neutralization were determined by comparing with the virus control. The data were analyzed and plotted with GraphPad Prism.

### Competition ELISA

Purified huACE2, CR3022, S309, P2S-2E9, BD55-5483, BD56-1854 and BD55-1205 were labeled with HRP (ProteinTech, PK20001). WT RBD was coated onto 96-well ELISA plates (50 ng/well). After blocking with blocking buffer (PBS with 10% FBS), blocking mAbs were added and incubated at 37°C for 1 hr. Then HRP-labeled detecting reagents were added in the presence of blocking mAbs and incubated at 37°C for 1 hr. The ratio of blocking and detecting reagents was 100:1. After wash, TMB substrate (Sangon Biotech, E661007-0100) was added and incubated at room temperature for 30 minutes. Then the reaction was stopped with stop solution and absorbance at 450 nm (OD450) was measured. The percentages of signal reduction caused by blocking mAbs were calculated.

### Binding activity measured by FACS

The plasmids encoding spikes of WT, BA.4/5, KP.3 and KP.3 mutants were transfected into HEK293T cells. After 48hr, cells were harvested. To compare the binding activity of germline and mature antibodies to WT and BA.4/5 spikes, cells were incubated with 3-fold serially diluted germline or mature mAbs (starting at 10 μg/ml) for 30 min at 4°C. To compare the binding with KP.3 and KP.3 mutants, cells were incubated with 1 μg/ml mAbs or 10 nM biotinylated ACE2 for 30 min at 4°C. After wash, cells stained with mAbs were further stained with DAPI and goat anti-human IgG FITC (Proteintech, SA00003-12), whereas cells stained with ACE2 were further stained with DAPI and streptavidin APC (Biolegend, 405243). After wash, cells were analyzed by CytoFLEX (BECKMAN COULTER). FACS data were analyzed with Flowjo V10 to determine the percentages of positive cells. To compare the binding activity of germline and mature antibodies to WT and BA.4/5 spikes, the percentages were ploted with GraphPad Prism. To compare the binding with KP.3 and KP.3 mutants, the percentages for KP.3 mutants were divided by the percentages for KP.3 and the ratios were recorded.

### Sequence analysis

Variable regions of antibody heavy and light chains from single B cells were analyzed with IgBlast (https://www.ncbi.nlm.nih.gov/igblast/). Using Python 3.12, clonotypes were analyzed with the Bio, pandas, and numpy packages. Antibodies with same V and J genes and more than 85% similarity in CDR3 (nucleotide sequence) for both heavy and light chains were defined as clones within the same clonotype. Spike sequences, uploaded from June 1 2024 to October 1 2024, were obtained from NCBI Virus. Sequences with lengths outside the range of 1250–1290 or containing ‘X’ were removed. Sequence alignment was performed using Muscle5, and statistical analysis was conducted with Python 3.12.

### Statistics and reproducibility

All statistical analyses were performed with GraphPad Prism. The numbers of biological repeats for each experiment and tests for statistical significance are described in corresponding figure legends.

## Supplemental material

Figures S1–S7

Data S1. Excel spreadsheet containing numerical data for figures.

## Data Availability Statement

All relevant data are within the manuscript and its supplemental files.

## Acknowledgements

The authors thank the donors for providing peripheral blood. The authors thank Professor Linqi Zhang and Peng Chen at Tsinghua University for confirming the pan-sarbecovirus neutralization of IGHV3-74 mAbs, and for providing plasmids encoding spikes of Pangolin CoV GD, Bat CoV WIV16, Bat CoV RaTG13 and plasmids encoding the heavy and light chains of mAb P2S-2E9. This work is supported by the National Key Plan for Scientific Research and Development of China (Grant No. 2022YFC2305800) and the National Natural Science Foundation of China (Grant No. 32200765, Grant No. 32370941).

## Author contributions

Conceptualization, L.J. and T.Z.; Validation, X.C., L.L., R.D., Z.W.; Formal Analysis, X.C., L.L., R.D., Z.W. and Y.L.; Investigation, X.C., L.L., R.D., Z.W., Y.L., Y.S., R.Q., H.F., L.H., X.C., M.L., X.H. and L.J.; Writing – Original Draft, X.C., L.L., R.D., Z.W., Y.L., L.J. and T.Z.; Writing – Review & Editing, L.J. and T.Z.; Visualization, X.C., L.L., Z.W. and Y.L.; Supervision, L.J. and T.Z.; Funding Acquisition, L.J. and T.Z..

---

**Figure S1.**
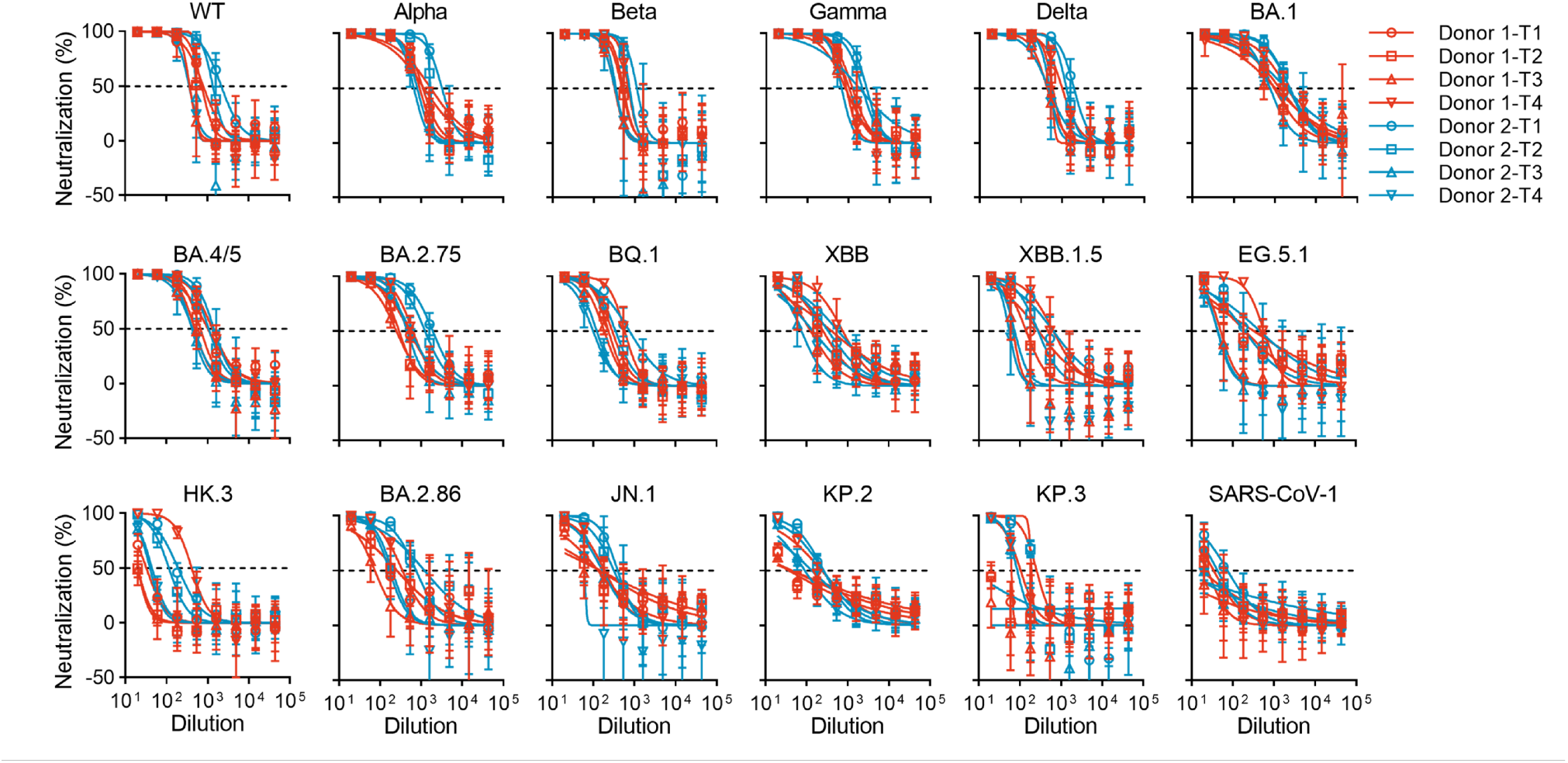
Plasma neutralizing antibody titers. (A) Plasma neutralizing antibody titers against a panel of pseudoviruses. The data are represented as mean ± SD and they are from three independent experiments, in which duplicates are performed. The published data for plasma from T1 are not used here. Instead, we measured the neutralizing tiers of plasma from T1 again, along with plasma from other time points.

**Figure S2.**
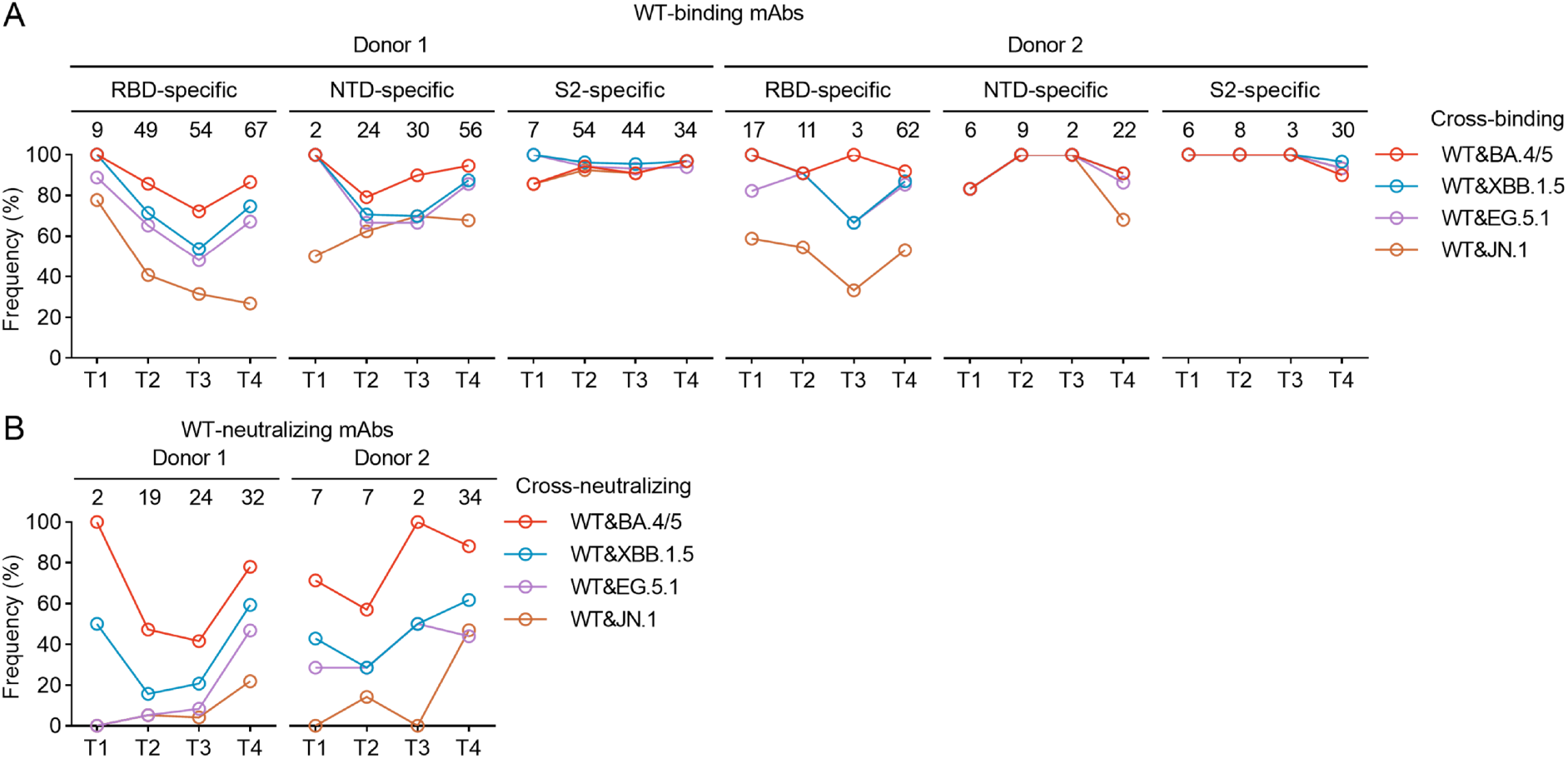
Cross-binding and cross-neutralizing mAbs from Donor 1 and Donor 2. (A) Frequencies of cross-binding mAbs among RBD-specific, NTD-specific, and S2-specific mAbs from Donor 1 and Donor 2. (B) Frequencies of cross-neutralizing mAbs from Donor 1 and Donor 2.

**Figure S3.**
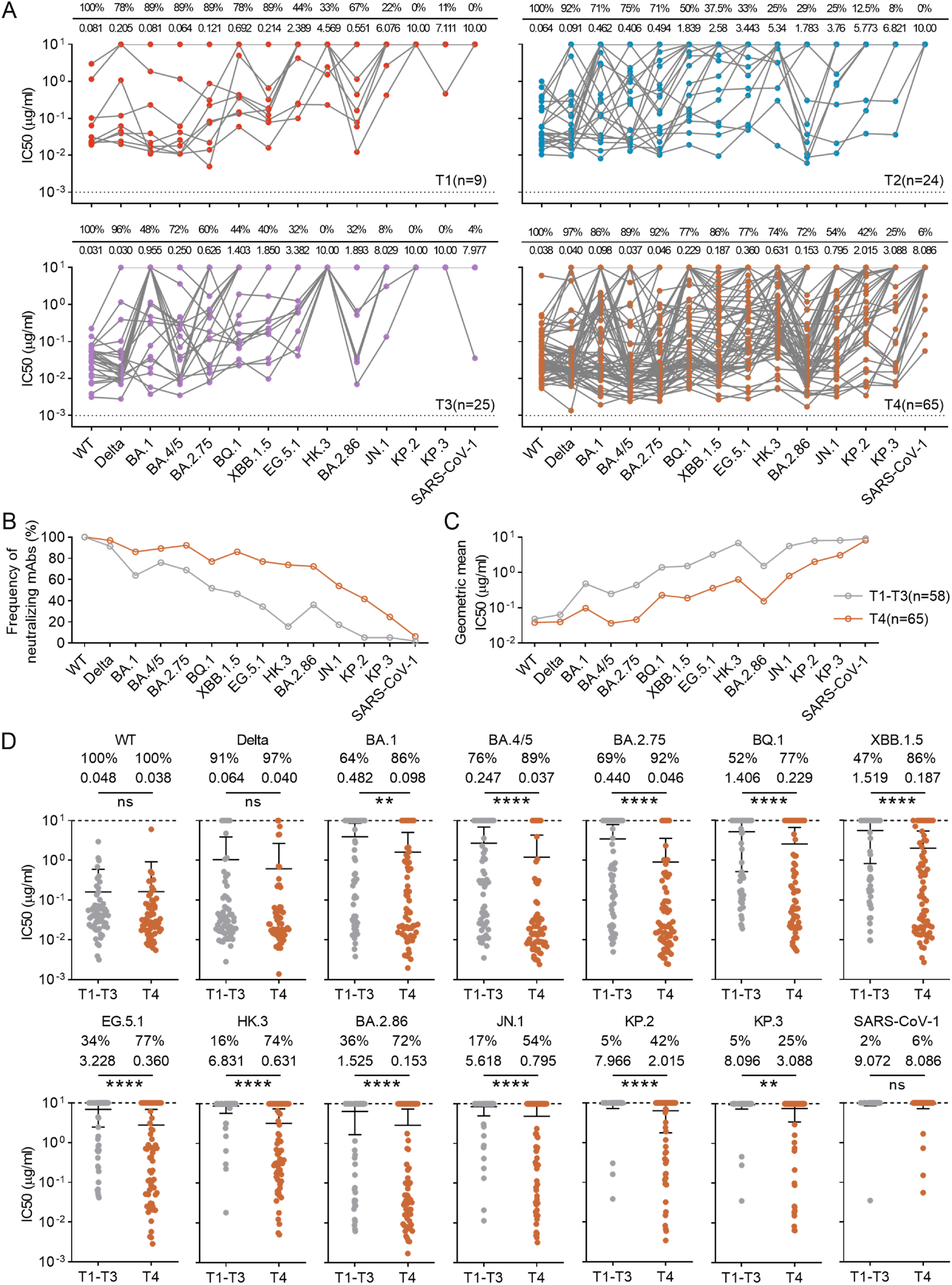
IC50s of mAbs. (A) Summary of IC50s. The numbers below the line indicate the geometric mean IC50s and the numbers above the line represent the percentage of mAbs that is able to neutralize the corresponding pseudoviruses (IC50 below 10 μg/ml). (B) Frequencies of neutralizing mAbs against each pseudoviruses among T1-T3 group and T4 group. (C) Geometric mean IC50s of mAbs among T1-T3 group and T4 group. (D) Comparison of IC50s. The numbers below and above indicate the geometric mean IC50s and the percentages of neutralizing mAbs, respectively. Statistical analysis is performed by Mann-Whitney test. ns (not significant) P > 0.05, **P < 0.01, ****P < 0.0001.

**Figure S4.**
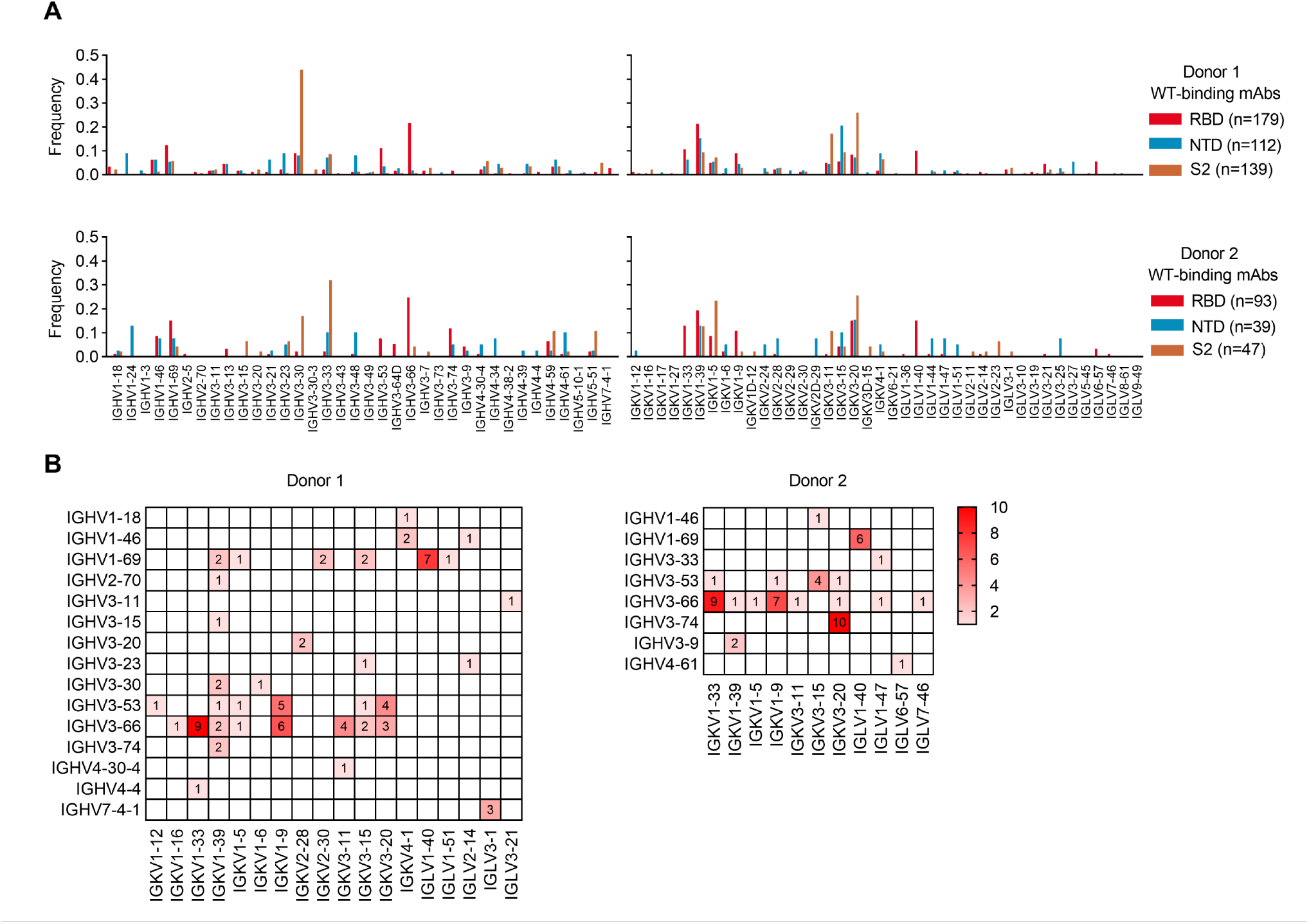
Analysis of V gene usage. (A) Frequencies of V genes used by mAbs targeting RBD, NTD and S2. (B) Paring of V genes for heavy and light chains in neutralizing mAbs. The number in each box indicates the count of mAbs using the corresponding pair.

**Figure S5.**
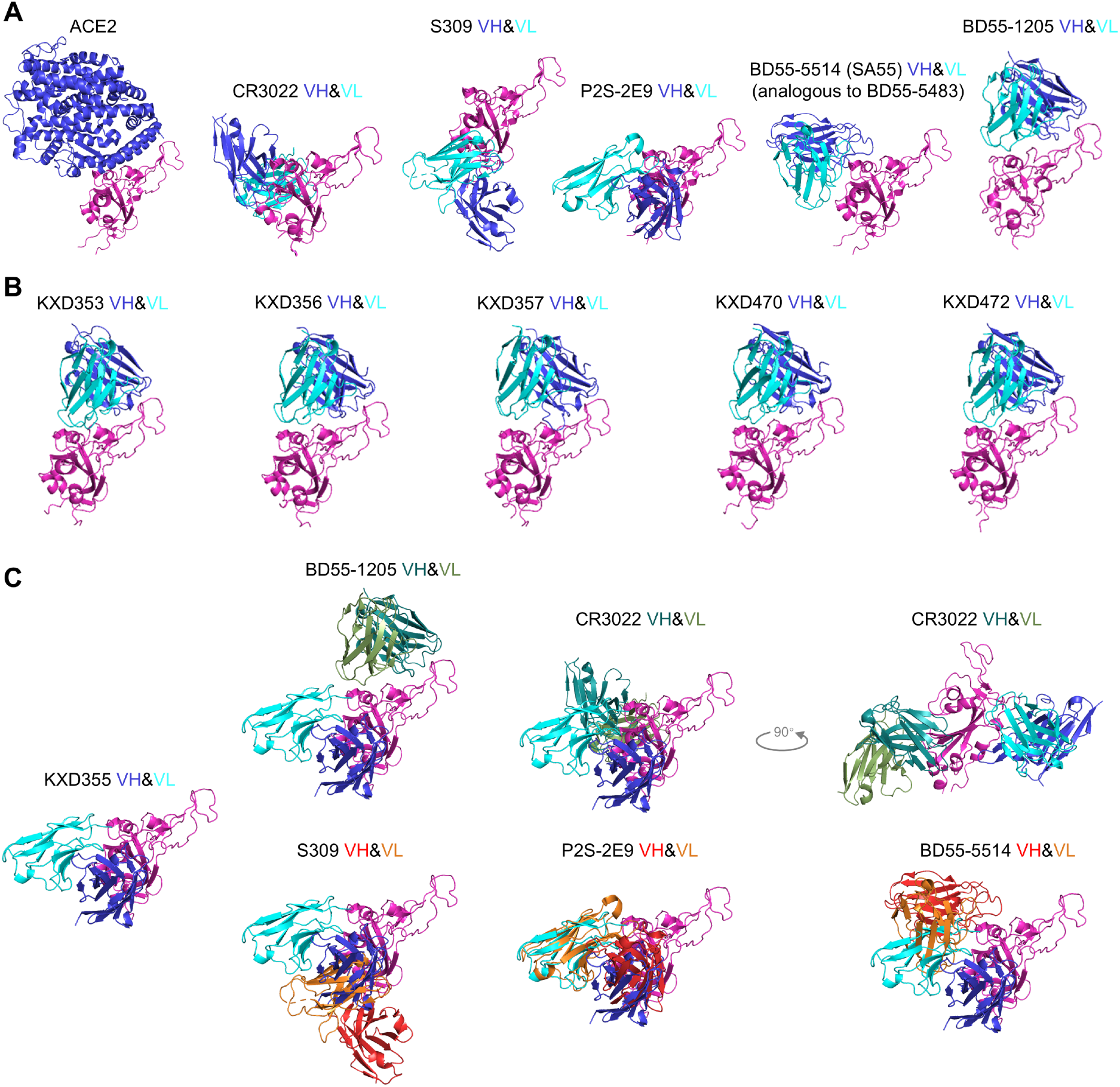
Published structures and predicted structures. (A) The published structures of ACE2 (PDB: 7A94), CR3022 (PDB: 6W41), S309 (PDB: 7TLY), P2S-2E9 (PDB: 7XSA), BD55-5514 (PBD: 7Y0W), BD55-1205 (PDB: 8XEA) in complex with RBD. (B) The predicted structures of KXD353, KXD356, KXD357, KXD470 and KXD472 in complex with WT RBD. (C) The predicted structure of KXD355 in complex with WT RBD is aligned with published structures of CR3022, S309, P2S-2E9, BD55-5514 and BD55-1205.

**Figure S6.**
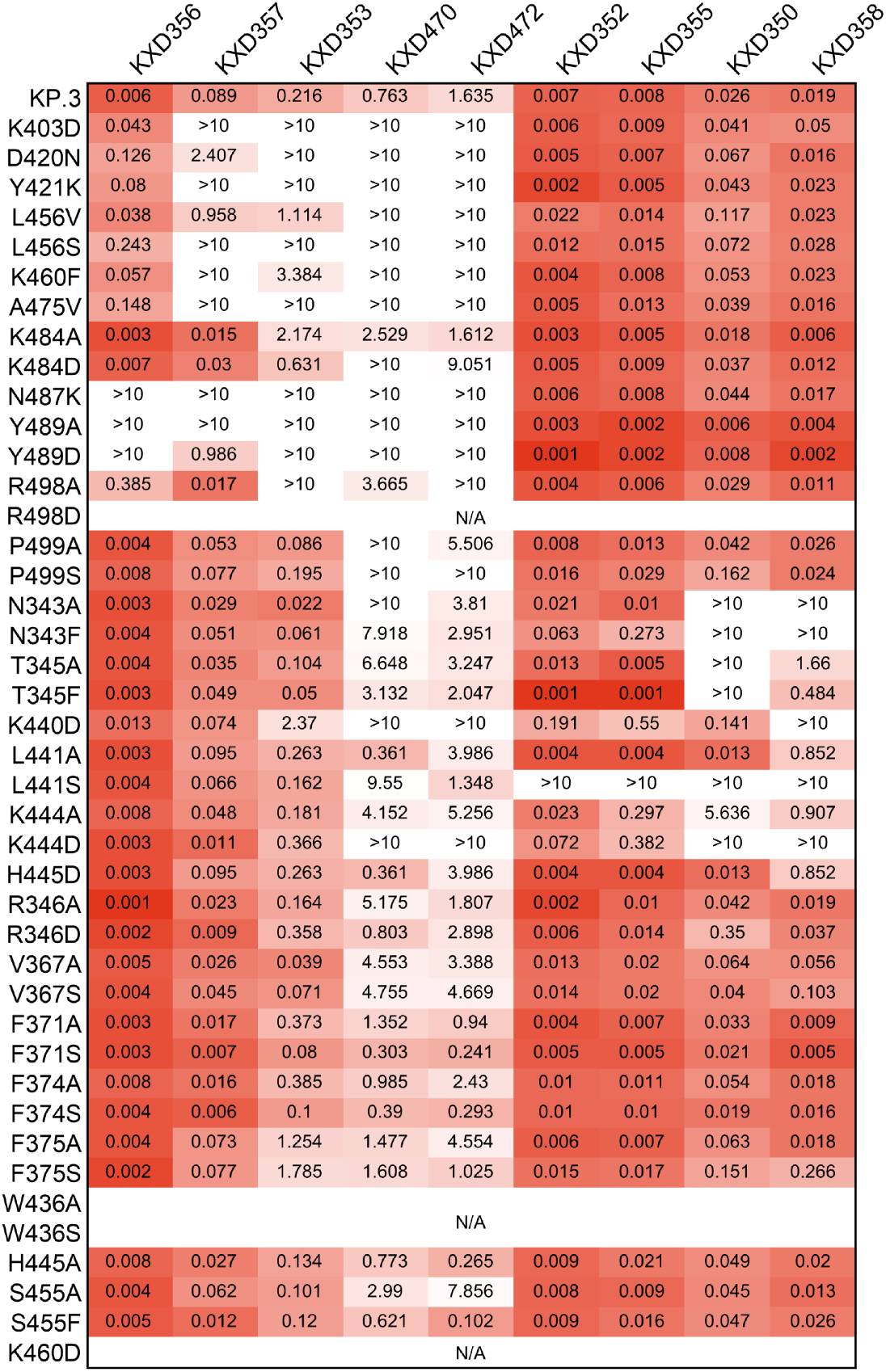
IC50s (μg/ml) against KP.3 mutants.

**Figure S7.**
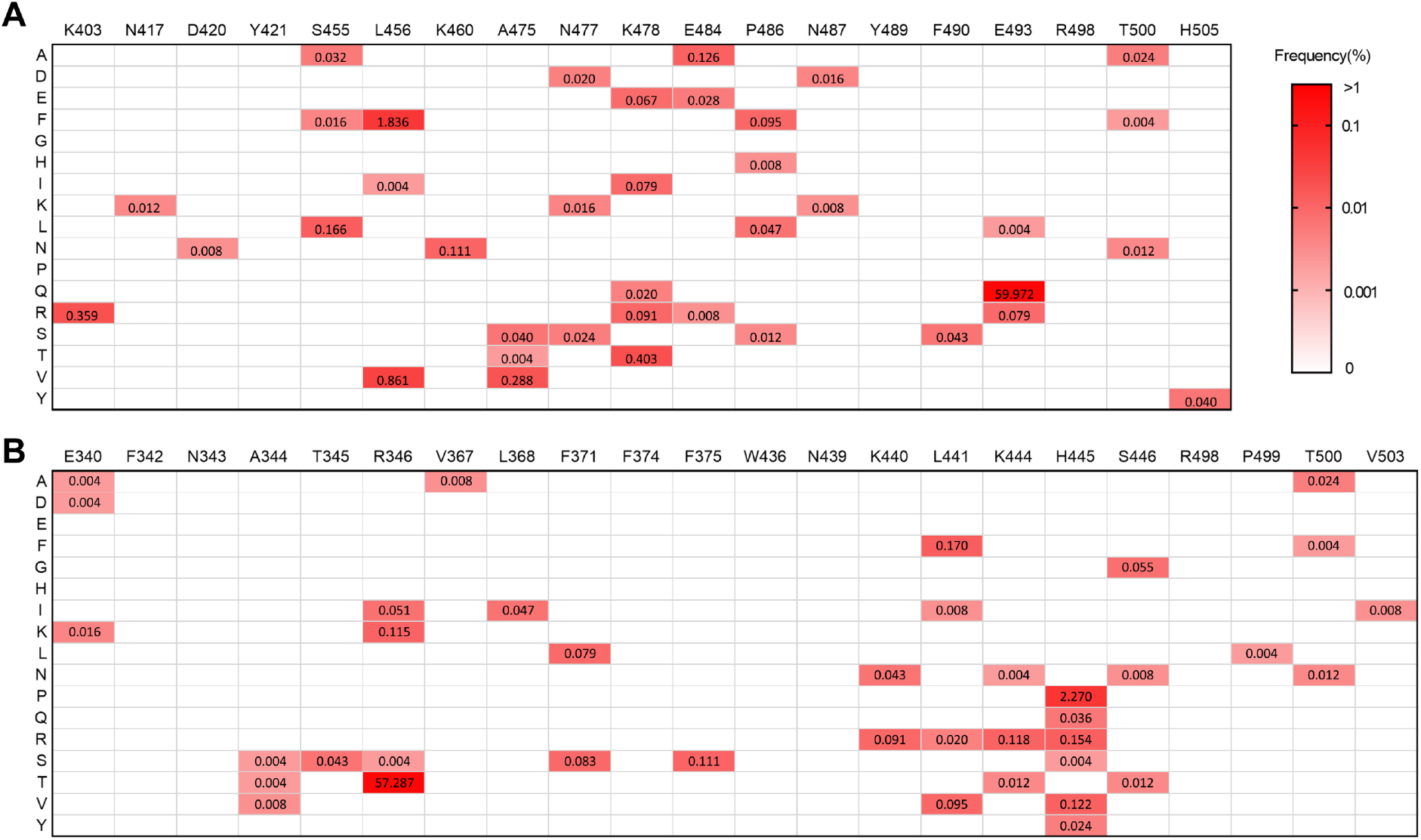
Variations on the epitopes of IGHV3-53/3-66 and IGHV3-74 mAbs. Frequencies of mutations on the epitopes of IGHV3-53/3-66 mAbs (A) and IGHV3-74 mAbs (B) among a total of 25332 spike sequences with KP.3 as reference. The sequences were published between June 1 to October 1 in 2024.

